## Supplementary information for "Selection, biophysical and structural analysis of synthetic nanobodies that effectively neutralize SARS-CoV-2"

**Supplementary Table 1 – Sequences of unique binders**

(provided as an excel file)

**Supplementary Fig. 1**

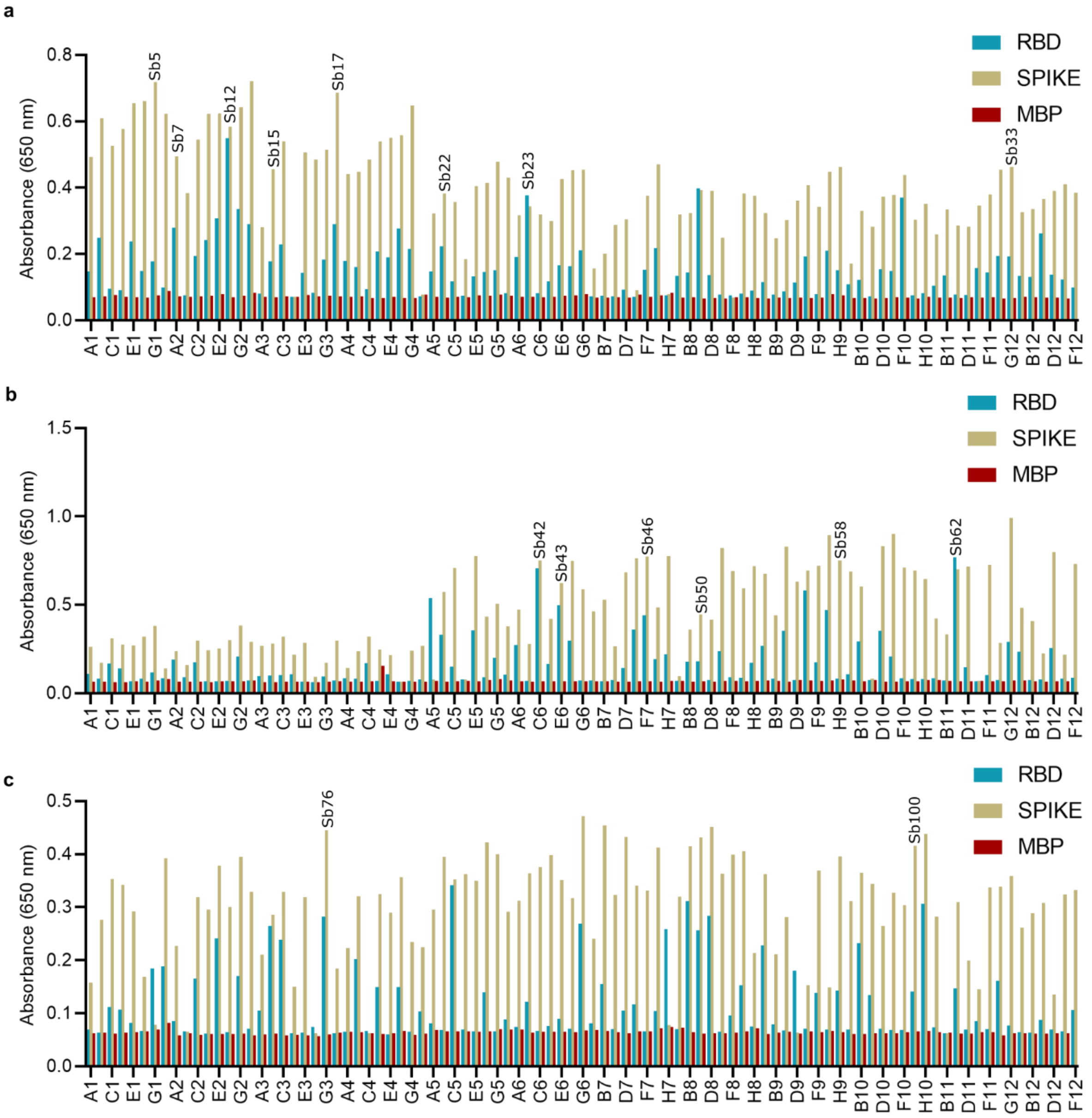


**RBD-specific sybodies were determined by ELISA.**

94 individual colonies from each of the three libraries, concave **(a)**, loop **(b)** and convex **(c)** were randomly selected and screened against the two target proteins, RBD and spike. The same procedure was performed for the MBP, used as background control signal. RBD or spike signals with ratios above 1.5 compared to the MBP signal, were considered as hits. The corresponding ELISA signals are labelled according to the sybodies that were illustrated or characterized in this work.

**Supplementary Fig. 2**

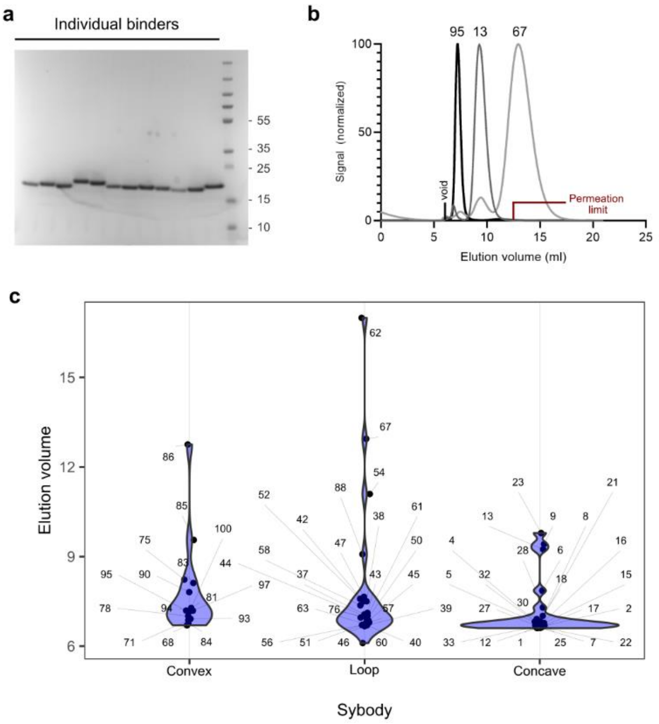


**Expression and Purification of 62 unique sybodies.**

From 85 unique binders, 62 were expressed and purified. The purified sybodies were analysed by SDS-PAGE **(a)** and gel filtration using a SRT SEC-100 column **(b)**. Very few sybodies exhibit column interaction (sticky binders), eluting at or after the permeation limit of columns. (**c**) Elution volume of all purified binders, grouped by library.

**Supplementary Fig. 3**

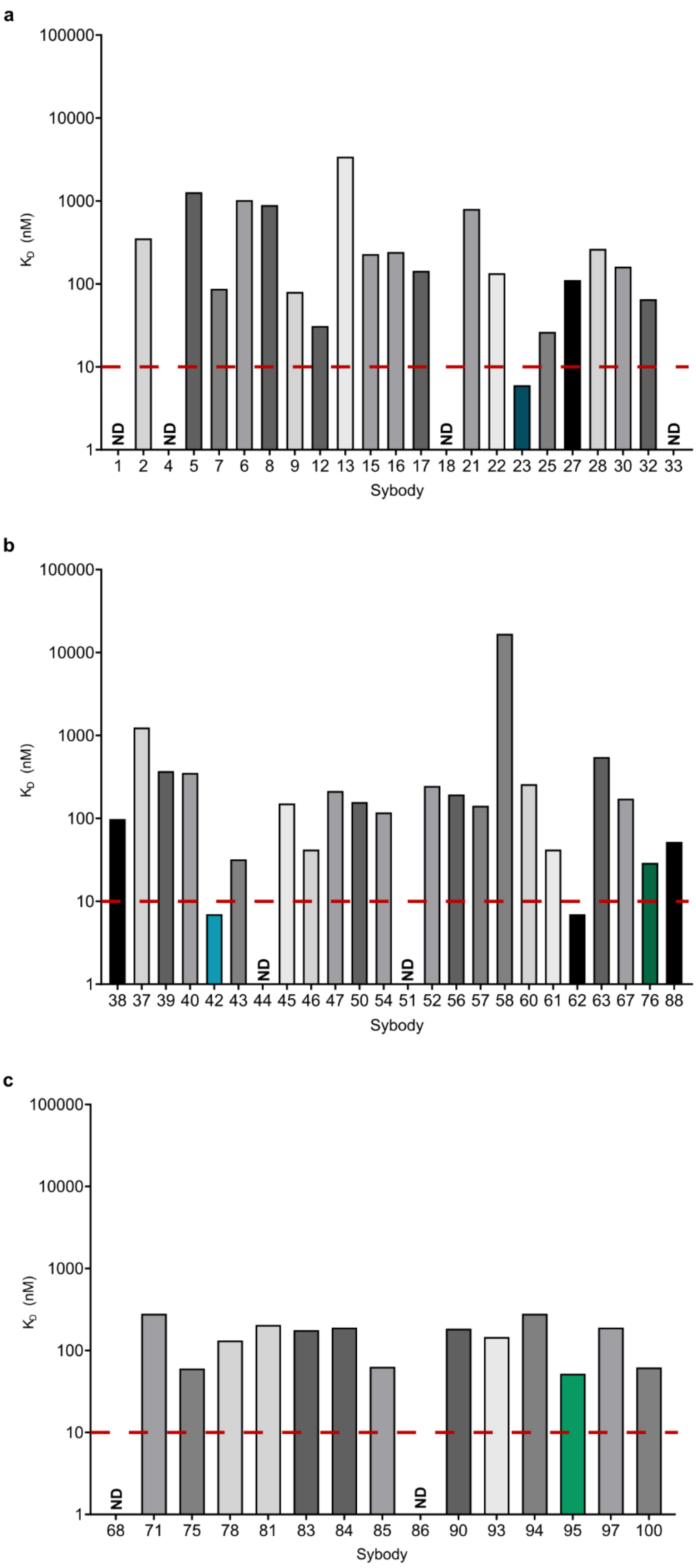

**Affinity screening of 62 sybodies.**

BLI sensorgrams of immobilized SARS-CoV-2 RBD with individual sybodies were recorded at one concentration (500 nM). The binding curves were fitted to a 1:1 binding model and $K_D$ values were estimated and plotted according to the different libraries concave **(a)**, loop **(b)** and convex (**c**). ND, not determined. The most promising binders, further characterized in this study, are highlighted.

**Supplementary Fig. 4**

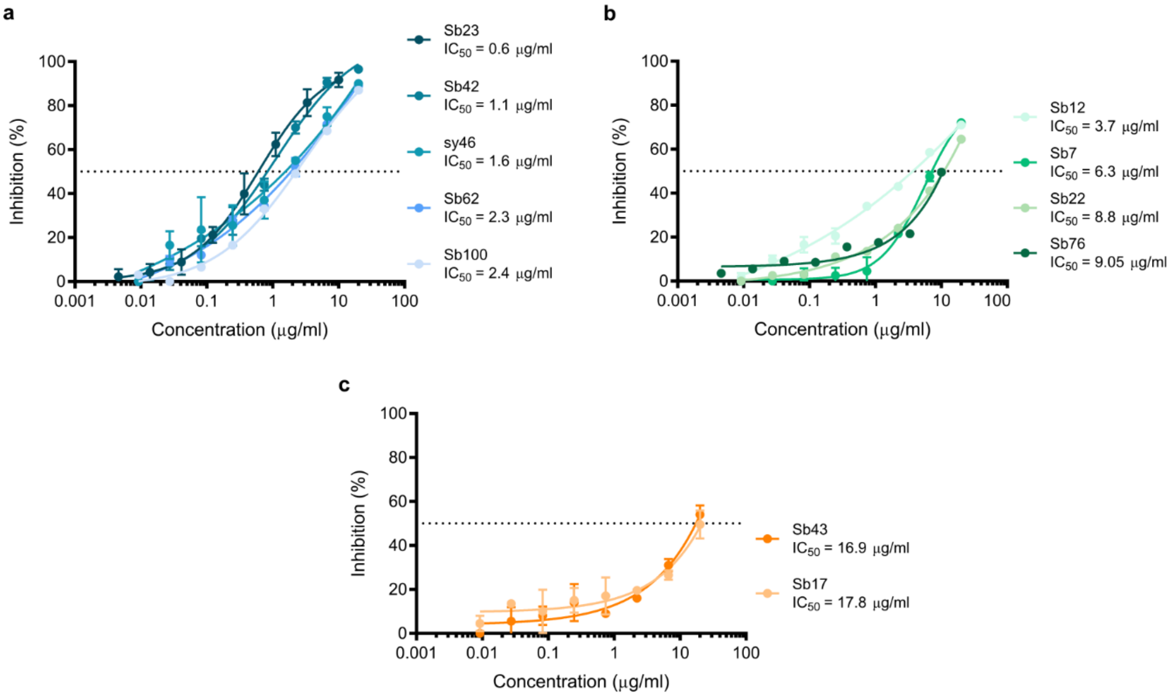


**Neutralization capacity of selected sybodies.**

SARS-CoV-2 pseudoviruses were incubated with a dilution series of target sybody. **(a)** Sybodies that showed the lowest $IC_{50}$ values. **(b)** Sybodies that showed $IC_{50}$ values above 3 µg/ml. **(c)** Sybodies that showed $IC_{50}$ values above 10 µg/ml. Error bars represent the standard deviation of at least two repetitions.

### Supplementary Fig. 5

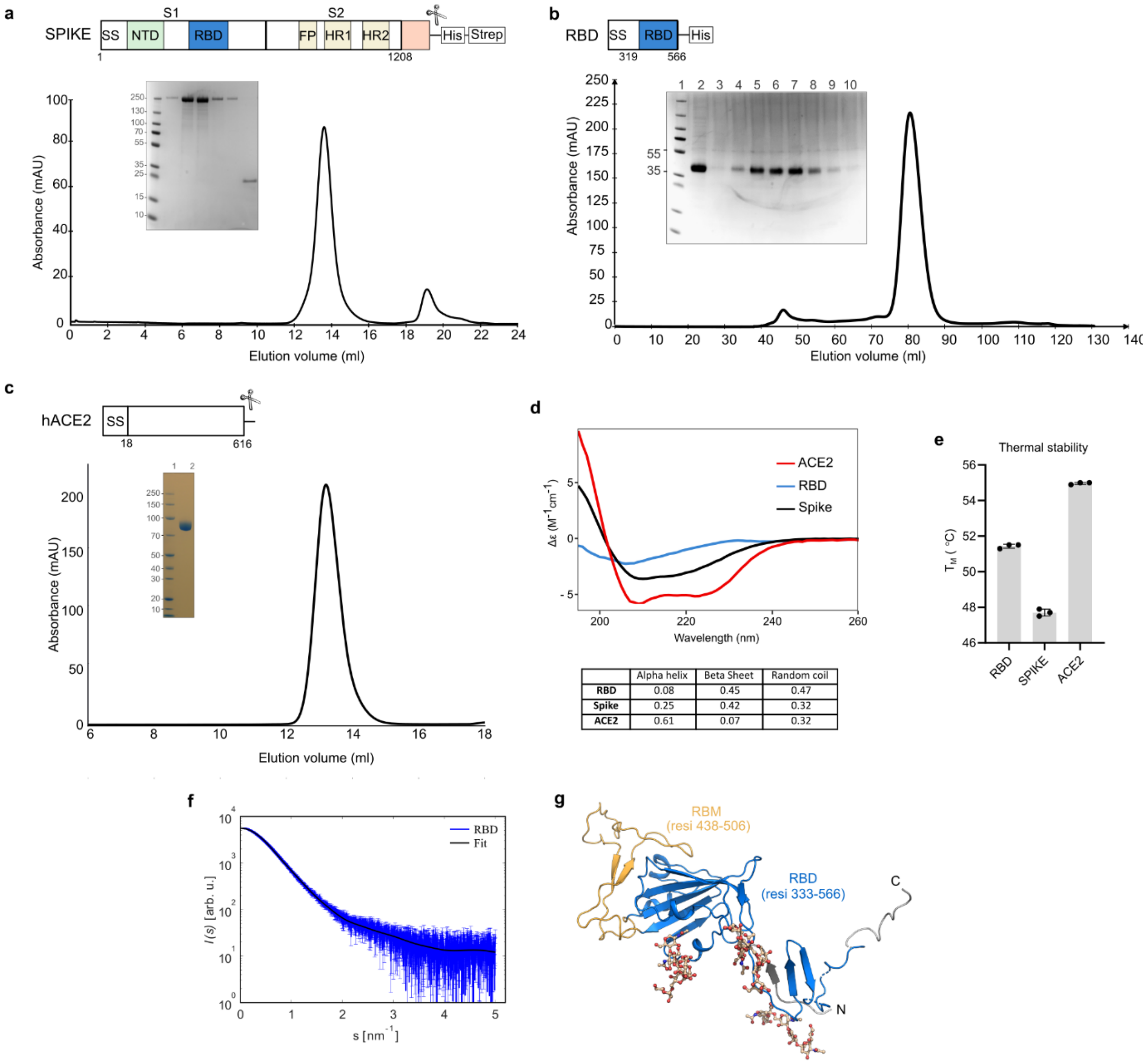


### Purification and biophysical characterization of antigens.

Purified proteins showed no signs of aggregation and remained stable and monodisperse. **(a)** Schematic illustration of the SARS-CoV-2 spike protein. Subunit S1 comprises the N-terminal signal sequence (SS), the N-terminal domain (NTD) and the Receptor Binding Domain (RBD), while subunit S2 consists of the fusion peptide (FP) and two heptad repeats, HR1 and HR2. At the C-terminus the construct contains a T4 fibritin trimerization motif, followed by a HRV3C protease cleavage site, an 8×Histidine tag and a Twin-Strep tag. Representative gel-filtration chromatogram of the spike protein. Inset: Instant blue-stained SDS-PAGE analysis of the size exclusion chromatography run. Lane 1: protein molecular weight marker. Lanes 2 - 6: fractions eluted from the Superose 6 column. Line 7: fractions contained 3C protease. **(b)** The RBD comprising residues 319 - 566 was used for sybody selections. A representative gel-filtration chromatogram of RBD is shown. Inset: Instant blue-stained SDS-PAGE analysis of the size exclusion chromatography run. Lane 1: protein molecular weight marker. Line 2: pooled fractions from IMAC purification. Lanes 3 - 10: fractions eluted from the HiLoad Superdex 200 column. **(c)** The human receptor ACE2 comprising residues 18 - 616 was used for competition assays. A representative gel-filtration chromatogram of the ACE2 protein is shown. Inset: Instant blue-stained SDS-PAGE analysis of the size exclusion chromatography run. Lane 1: protein molecular weight marker. Lane 2 pooled fractions eluted from the Superdex 200 column. (**d)** Far-UV-CD spectra of the human ACE2 receptor (red), SARS-CoV-2 RBD (blue) and SARS-CoV-2 spike (black) with the corresponding secondary structure content. (**e)** Thermal stability analysis of RBD, spike and ACE2. (**f)** Experimental SAXS data from RBD and the computed fit from the hybrid model of RBD displayed in g. The SAXS profiles are presented as intensity in logarithmic scale versus momentum transfer $s = (4\pi \sin\theta)/\lambda$, where $2\theta$ is the scattering angle and $\lambda$ is the X-ray wavelength. **(g)** Cartoon representation of the corresponding RBD model. The RBD is coloured in blue and the receptor binding motif (RBM) is coloured in orange.
